# Reactions to Perceiving Tangible vs Virtual Impossible objects: Dimensions of Visual Reality

**DOI:** 10.64898/2026.10.14.759059

**Authors:** M. Souilhol, A. Foncelle, J. El Khoury, A. Seye, F. Volland, F. Quesque, A. Kambara, Y. Rossetti

## Abstract

Considering the current plethora of 2D depictions via flat screens, we investigated whether our perception might have become more susceptible to this format or whether our perceptual system remained more captivated by tangible reality, i.e. 3D objects. To this end, we selected a set of impossible optical illusions that can be described as impossible objects and evaluated their impact on perceptual experience. Twenty-eight healthy volunteers were presented with 5 impossible objects in three ways: real wooden format, photographs or coloured drawings of these objects. They had to evaluate their reactions to each illusory stimulus separately in a random order. Then they were asked to rank the three presentations when presented simultaneously. We found that real objects presentation was more challenging than their 2D counterparts. Specifically real objects yield more surprise and impact on perception than their 2D image counterparts. Implications for naïve realism and social cognition are proposed.

## Introduction

During one of the very first showing of a film (Lyon, France: *Arrival of a Train at La Ciotat Station^1^*), screened by the Lumière Brothers, 1896 – the legend says that everyone in the audience took leave of their senses and some broke into panic and ran for their lives. Actually, perceiving our environment in three dimensions (3D) seems easier and more transparent to us than breathing. Yet, although we are accustomed to perceiving and moving through a 3D environment, most of the information and signals we receive are actually two-dimensional (2D).Our visual perception is mediated by our retinal receptors, which capture 2D projections. Our organisms have learned to efficiently decode these signals in order to understand and interact harmoniously with the 3D physical world. This ability allows us to cooperate with living beings, avoid dangers, and find partners or food, thereby ensuring our survival and that of the species.

Although this mechanism of translating 2D signals into coherent 3D percepts seems both trivial and elementary, since perception feels so natural to us, it involves numerous and complex mechanisms for processing and transforming 2D information. As Welchman (2016) points out, our perceptual system flexibly and robustly apprehends the depth dimension from a variety of 2D stimuli, without us even paying attention to it. As a matter of facts, depth perception can be derived from a number of both binocular and monocular cues (Tsutsui et al. 2006; Sugihara 1986; Welchman 2016; Snow and Culham 2021). These cues depend both on retinal signals and higher-level processing (Trotter 1995; Bridge 2016; Bridge et al. 2023). Among the parameters contributing to depth perception, perspective, parallax, motion, rectangularity, and binocularity play particularly central roles (Glennerster, 2016; Sugihara & Pinna, 2024; Li et al. 2025), allowing the visual system to construct robust representations of 3D space. Thus, many factors come into play in translating 2D information into 3D percepts within the context of the deceptively simple visual perception.

Historically, one way to highlight the complexity of the operations required to convert 2D information into a 3D environment is to rely on illusions, i.e. on perceptual phenomena in which the sensory experience differs from the physical reality of the stimulus. Indeed, since at least Aristotle, visual illusions have been presented as puzzles whose solutions help to better understand not only the workings of perception but also the nature of reality. Understanding the errors in translation that we make when perceiving certain objects allows us to better grasp the operations we perform unconsciously when perceiving our environment. Becoming aware of these operations encourages us to take a step back from what we think is reality. It leads us to question our naive realism, that is, the natural conviction we form that we perceive things exactly as they are. For instance, impossible objects (e.g., the Reutersvärd-Penrose triangle, Fukuda’s disappearing column) primarily engage rectangularity processing (Sugihara and Pinna, 2024) and lead to altered visual processing (e.g., Shigemura et al., 2004), thereby allowing the study of the mechanisms underlying visual perception and challenging our understanding of what is materially possible or impossible.

Encapsulated three-dimensional interpretations of 2-D information leads to a paradox: on the one hand, it is long known that presumably most striking visual illusions rely on the fact that our perceptual system has developed sophisticated strategies to infer the 3D from 2D retinal projections of the world (Shapiro and Todorovic, 2017) but, on the other hand, visual illusions have long been largely limited to 2D materialization of 3D objects. Indeed, in most research on visual perception and illusions, the stimuli used are essentially 2D (Linton, 2016; Ferretti, 2024; Snow and Culham, 2021). Additionally, experimental approaches to illusions have remained restricted to 2D images even in the case of studies of visually guided actions and interactions with the external world (e.g. Coello et al. 2007). Whether they are drawings, photographs, geometric figures, or videos, these flat representations are easy to present and control experimentally. They allow precise study of perceptual mechanisms, misinterpretations, and illusion effects, but they remain limited because they do not fully reproduce the 3D complexity of the real environment in which we operate (Snow and Culham, 2021). Our technological evolution is such that we are now constantly exposed to a continuous stream of 2D images, much more than a hundred years ago at the time of Lumière brothers. Between television, computer, and smartphone screens, as well as social media, our visual environment is largely dominated by flat representations of the world. Today, the average world individual spends as much as 6 hours and 40 minutes daily on a screen, with a peak value around 9 hours daily for the generation Z (datareportal.com, 5 February 2025), which exceeds half of the daily waking hours! This intense exposure may subsequently alter the perceptual status of 2D and 3D stimuli, enhancing our ability to interpret 2D cues to mentally reconstruct depth, while perhaps also altering the way we perceive and interact with our real 3D environment. Also, the rapid digital dissemination of information has made it increasingly challenging to determine what is real and veridical, so much so that fake and deepfake have recently become a serious educational issue (Peters, 2017).

There are a few historical exceptions to this paradox in the scientific domain (e.g., Beuchet’s chair (1963, see Devinck and Quaireau 2024), von Helmholtz & Ames’ room, 1946), while there has been a flourishing production of 3D objects in the artistic domain, ranging from street art to renowned names (e.g. Salvador Dali, Maurits Cornelis Escher, Shigeo Fukuda, Oscar Reutersvärd, Jiro Tani, Victor Vasarely,…). Thus, material 3D implementations of the 2D concepts have long remained the privilege of artists^2^. It is time for research on visual perception to build on these technical advances to continue questioning our understanding of vision and reality.

The 3D implementations of physically built impossible objects were initially derived from preexisting 2D graphical models. One of the main properties of those objects is that they may well be designed in 2D (e.g. Dali, Escher, Fukuda, Magritte or Reutersvärd) but it is not feasible, by definition, to translate them into tangible 3D objects. Thus, materializing impossible objects is inherently challenging due to their paradoxical composition. Because of this challenge, 3D materialization of impossible objects or ambiguous objects are seldom encountered as compared to 2D (e.g. J Beuchet’s “cube-woman”, M Rätz’ untitled “oui-non”, 2000; G Moretti, “Cubosfera” series 2011). In addition, because sharing information is usually achieved by 2D medias (classical papers or modern screens), the intrinsic value of their tangibility is somehow degraded, and they paradoxically do not overtake the success of their 2D predecessors. It should be emphasized that a photograph or drawing of an object is already a representation, i.e. an artifact for which we have been used to deal with distance from reality (Ferretti 2016). As Magritte emphasized, we cannot smoke his painting of a pipe (Magritte’s “La trahison des images (The Treachery of Images), 1929). This distance should even more be increased for impossible objects. By contrast, the physical presence of a tangible wooden object should be expected to create a stronger cognitive conflict. The object is there, materialy present, and yet it cannot exist. This direct confrontation with an “impossible reality” should generate a deeper surprise and doubt than the observation of a mere image of it.

Through our own teaching and dissemination experience we gradually developed the conviction that being confronted to materialized illusions (e.g. size-weight illusion) or to wooden-made impossible objects was more challenging to perception that the most-spread 2D illusions. Thus, the case of impossible objects (Sugihara 1982) is extremely interesting to consider because they effectively trigger surprise, puzzlement, and metacognition and therefore can be used to uniquely address issues ranging from philosophy, perception, reality, aesthetics, neuroscience and cognition as they can (Bach and Poloschek 2006; Krause et al. 2019; Quesque et al. 2023). We hypothesize that these impossible objects therefore allow not only the study of perceptual mechanisms but also the questioning of our understanding of perception and reality, i.e. the process of interrogating apparent evidence and assumptions (Dokic and Martin 2017; Textor 2019).

The Achilles’ heel of these impossible objects is that they have to been seen from a single accidental view, and, hence, that free viewing will immediately reveal that they are actually not impossible objects (Li et al. 2025). In order to prevent the observer from uncovering the forced perspective effects, they should be given a very restricted visual access, i.e. using monocular presentation through a pinhole so that the fragile illusory effect doesn’t collapse (Sugihara 2020). One interesting aspect within the recent screen (r)evolution is the sustained enthusiasm for discovering and sharing new visual illusions. Numerous websites and blogs display not only classical 2D illusions but more interestingly images of the 3D world, e.g. forced perspective showing a hand grasping the sun or someone standing on someone else hand. Therefore, the two-dimensional format allows for ideal presentation of illusory scenes or devices that would not appear as saliently if there were to be observed as real three-dimensional scenes as our two eyes and minimal head or body motion allow for disambiguating spatial alignments, forced perspective and depth confusion.

The first goal of this study was to investigate whether three-dimensional stimuli may actually induce more powerful effects on the perceptual system, even if the viewing angle can obviously not be as narrowly constrained as for flat images. As illusions have been recently proposed to be a promising tool to develop openness to the other’s point of view (Kambara 2017, 2021; Quesque et al. 2023), our secondary question was about whether experiencing the limitation of a single spatial perspective via exposure to impossible objects may also transfer to cognitive perspective taking.

## Material and methods

Twenty-eight healthy volunteers (11 females, 17 males; age range: 19–66 years) participated in the experiment and provided written informed consent in accordance with the Declaration of Helsinki. No additional ethical approval was sought, as the study involved only non-invasive behavioral measures with adult volunteers. The sample size of N = 28 was chosen to approximately double the number of participants in an experiment using similar material (Sugihara & Pinna, 2024), ensuring sufficient power for repeated-measures comparisons. A post-hoc power analysis confirmed that this sample provides adequate power to detect medium-to-large effects (Cohen’s f = 0.25–0.40, see Annex 1).

Five classic optical illusions were selected as stimuli: four impossible objects (cube, Penrose triangle, impossible frame, Blivet) and one cognitive illusion (Oppel-Kundt). Each illusion was presented in three formats—real wooden object, color photograph, and line drawing— yielding a total of 15 stimuli (see Figure 1, Table 1). Lighting and viewing conditions were controlled to maximize the perceptual impact of each illusion.

**Figure 1.**
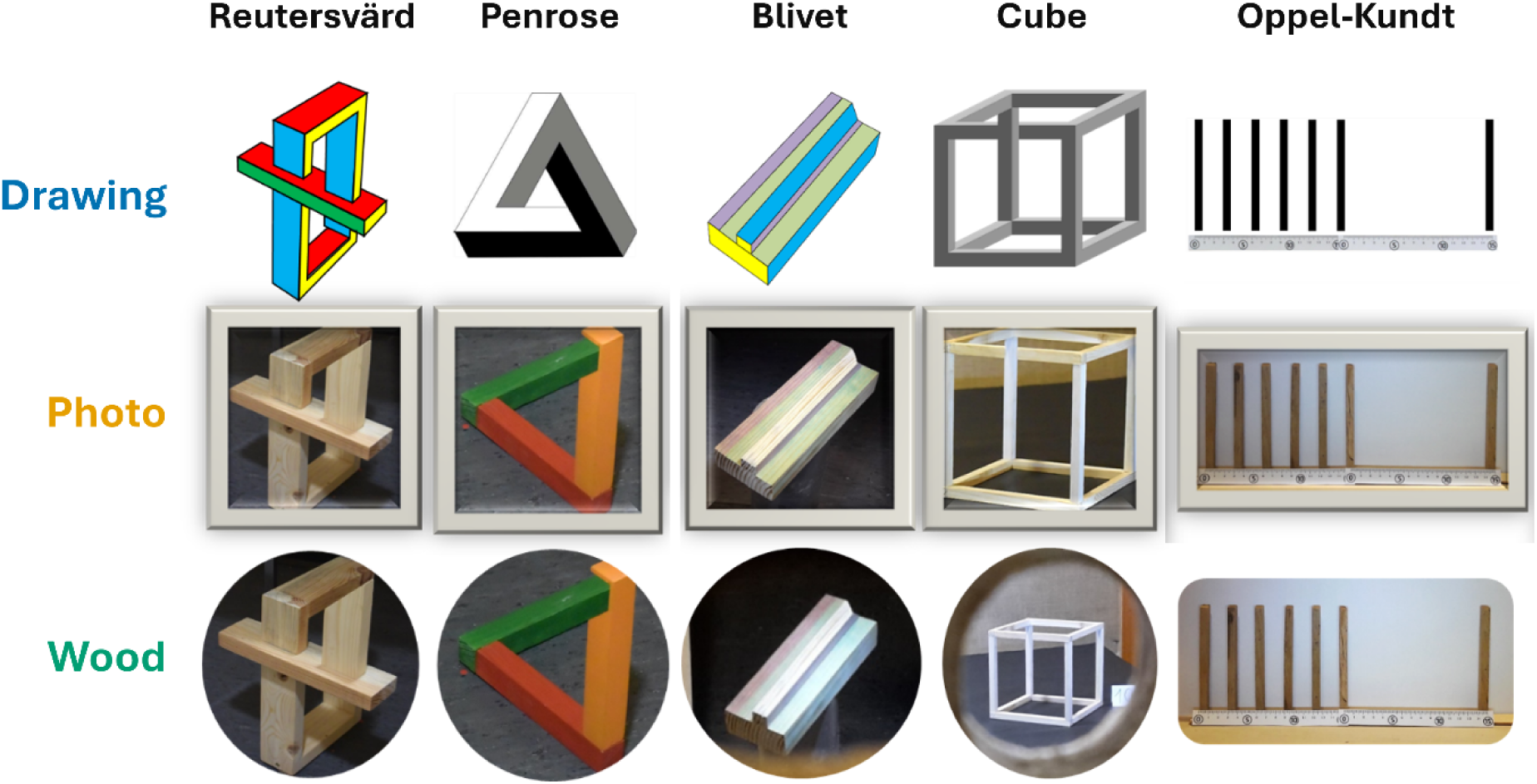
The five illusions and their three presentations. We first selected 4 strikingly impossible objects and one more cognitive illusion. From left to right the figure depicts an impossible frame (Oscar Reutersvärd, “Perspective Japonaise” series) and created in 3D by Hitoshi Ikeuchi (2005; https://thanks.yahoo.co.jp/ikemath/album/illusion.html), the Penrose-Reutersvärd triangle (which will be unjustly^5^ and simply referred to as the Penrose triangle, as another illusion will be referred to as the Reutersvärd illusion here; Mortensen 2022, Sugihara 2021), a Blivet (Bach & Poloschek 2006; also designed by Oscar Reutersvärd, and an impossible cube (a version of the Necker cube in which edges cross in inconsistent ways). The four impossible objects, Reutersvärd structure (L37xW30xH48cm), Penrose-Reutersvärd triangle (L23xW20xH23.5cm), the Blivet Block (L13.3xW7.8xH3.7cm), a cube (16.5cm), and the Oppel-Kundt filled-space (L40xW3xH12.5cm) illusion were used as stimuli and presented as real wooden objects (bottom raw). In addition to these tangible objects, pictures of the real objects (middle raw), or drawings (top raw). Additionally, the Oppel-Kundt filled gap illusion was made in 3D with Kapla^®^ wooden blocks and printed rulers (right-most column). Rulers were added to the Oppel-Kundt display to enable participants to perceive its illusory dimension without explicit description. We thus used a total of 5×3 =15 different stimuli. Lighting of the 3D objects was carefully adjusted to enhance the optical illusion of each stimulus. For example, the backmost pillar of the impossible cube was specifically lit in order to enhance the feeling that it was actually in the front of the cube.

**Table 1:** The five illusions used.

| **Stimulus** | **Description** | **Perceptual conflict** |
| --- | --- | --- |
| Impossible frame<br>(Reutersvård-Ikeuchi frame) | Rectangular frame where an additional plank is added which appears to connect the left-right and the front-back planes in a geometrically impossible manner. | Makes the 3D interpretation impossible as the viewed object cannot be materialized in 3D Euclidian space. |
| Reutersvård-Penrose triangle | Impossible 3D-like triangle composed of three straight beams of square cross-section, which meets pairwise at right angles. | Creates a spatial aberration as it forms a closed loop that cannot exist in real 3D Euclidean space. |
| Blivet Block | Derived from the famous three-pronged trident-like figure known as the devil's Tuning Fork, the central element of this object deceptively transitions from being convex at one end to being concave on the opposite edge. | Creates a deceptively transitions from convexity to concavity, creating an irreconcilable conflict in depth apprehension and 3D reconstruction. |
| Impossible Cube (Necker-Escher cube) | Classical hollow cube whose structural bars intersect in contradictory ways in two crossings. | Forces the visual system to perceive background and foreground edges simultaneously in the same spatial plane. |
| Oppel-Kundt | Classic cognitive/geometrical illusion where a divided or filled space (usually packed with vertical ticks) is systematically overestimated and perceived as wider than a physically equal, unfilled empty space. | Violates geometry rules without implying perspective cues. Perceptual conflict arises here from the rulers allowing to compare the filled and unfilled spaces. |

### Procedure

Participants had no prior knowledge about the scientific aim of the study. All measures, manipulations, and exclusions are reported in the present report. The protocols were in accordance with the principles of the Declaration of Helsinki. The volunteer entered the experimental room and faced a wide wooden panel preventing them to any of the stimuli. Then they sat on a chair and placed one eye in front of a drilled hole (diameter 5mm), which could be occluded by the experimenter on the side of the stimuli. There was no window in the room so that the viewing conditions of the illusions were kept constant. All the optical illusions were placed under a table (110 to 150 cm from the viewing point) or above (depending on the appropriate viewing angle) and they remained hidden so that participants could only see one illusion at a time and focus on it. Volunteers freely watched the optical illusions and had no hint about what they had to guess or see.

Attention was paid to make the 2D and the 3D stimuli as comparable as possible: viewing distance and visual angle sustained by stimuli were similar, and no pixelation was visible on the images. Nevertheless, the perceptual difference between the picture and the 3D object was obvious to participants as they could view a large scene (approx. 30°) in which the object or the printed page were presented. In the real-world condition, observers had access to natural depth cues such as accommodation, depth of field, and micro-motion parallax. Although our ecological approach did not aim to isolate the individual contribution of each of these optical cues, they may interact with higher-level cognitive processes. Crucially, the major distinction between the two formats lies in the perception of the boundaries of the medium. In the photographed condition, the visible edges of the display or print serve as a powerful cognitive cue that the stimulus is merely a representation. As emphasized by Ferretti (2024), viewing the edge of an image is a crucial cue to distinguishing between reality and images and trompe-l’oeil stimuli do not lead to illusions as long as their frame is visible. In contrast, the seamless integration of the impossible object within the viewer’s actual physical environment in the wooden objects condition lacks these restrictive boundaries, thereby triggering a genuine expectation of physical reality and heightening the subsequent cognitive surprise. In addition, the viewing-hole size was sufficient to enable slight motion parallax and active exploration by eye movements, explaining why the Blivet illusion induced smaller difference between 2D and 3D presentations (Supplementary figure A2).

The experimental phase went through two steps. The first step consisted in an implicit rating of each stimulus (i.e. 5 illusions x 3 presentations = 15 items). Volunteers freely viewed each of the 15 stimuli and then rated them on 10 cm analogical scales depicting “not at all” on their lefthand side and “extremely” on their righthand side. The following questions were asked Q1: how much they feel surprised, Q2: how much they feel disconcerted, Q3: how much they rate their curiosity for this stimulus and Q4: at which point the stimulus makes them doubt about the objectivity of their perception, i.e. the capacity to perceive an object as it exists independently of the perceiving subject. This allowed us to compute average numerical values from which an implicit ranking was derived. During the second step, an explicit ranking was obtained from each participant during the simultaneous presentation of the three versions of each illusion (i.e. 5 trials). Participants were asked to rank them (1 to 3) along three dimensions: surprised, disconcerted, and doubting, i.e. estimating the perceptual challenge imposed on the visual system by these stimuli. Surprise serves as a direct behavioral indicator of a violation of internal predictive models when facing structurally anomalous objects. The order of presentation of the stimuli was pseudo-randomized, such that they would never see the same stimulus (i.e. 5 illusory set-ups) or the same type of presentation (3D wooden object, 2D photographs or drawing) in consecutive trials.

## Results

### Qualitative observations

Individual sessions lasted about 35 minutes. Participants spontaneously commented that they enjoyed the experiment and did not encounter difficulties. Although we could not see participant’s face during the experimental sessions, it is worth mentioning that they spontaneously expressed surprise by vocal or verbal reactions. This happened above all for the Wooden presentation. Additionally, 11 out of 28 participants provided comments on their difficulty to ‘understand’ the Oppel-Kundt stimulus and apprehend the illusion.

### Scoring analysis

The scores provided by each subject (n=28) for each illusion (n=5) and each version (n=3 presentations), and for each question (n=4) were subjected to a repeated measure analysis of variance. All ANOVA assumptions were checked.

The **Illusion** main effect was significant (F(4,108) = 16.5; P<0.001, η²p=0.379), and mainly driven by the fact that the Oppel-Kundt Illusion yielded smaller scores than the others four (see Annexes figure A1; all ts>4.0; Cohen’s ds>0.80; holm adjusted Ps<0.003) (see **figure 2**).

**Figure 2.**
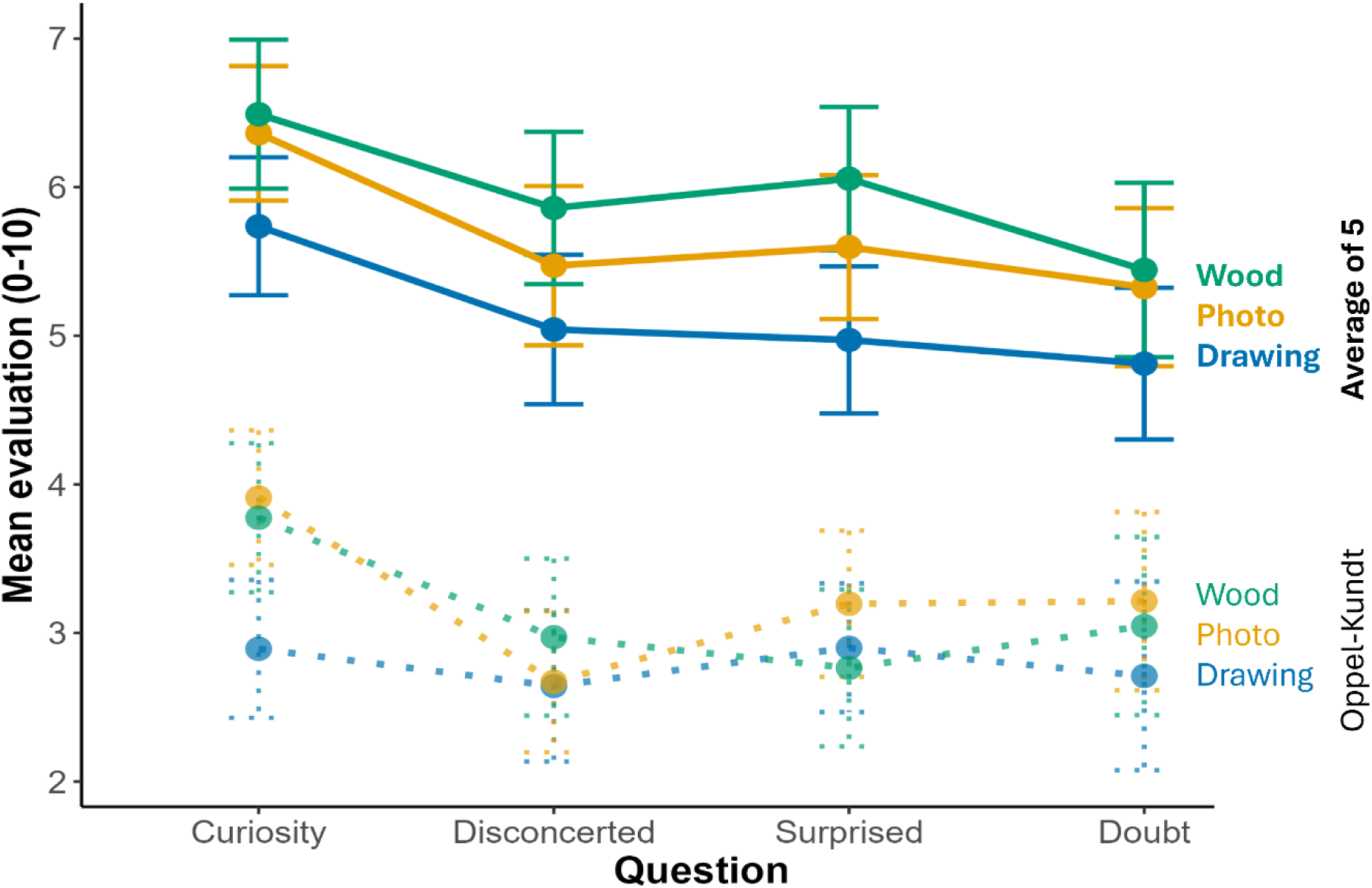
Illusion types and versions effect on cognitive evaluations. Mean scores (±SE) showed that the Drawing version was the least effective, whereas the Wooden version was the most effective for impossible object stimuli. Oppel-Kundt illusion significantly departed from the other illusions and is represented separately here. The four other specific illusion drawings can be found in Annexes. Evaluation scores ranged from 0 (no effect) to 10 (maximal effect). Error bars depict standard errors of the mean.

In addition, the Reutersvärd illusion obtained significantly higher scores than the Cube four (t=2.97; Cohen’s d=0.34; holm adjusted P=0.037) and than the Penrose triangle (t=2.84; Cohen’s d=0.33; holm adjusted P=0.043). The significant **Version** main effect (F(2,54) = 7.00; P<0.002; η²p=0.206) was mainly supported by the fact that the Drawing version was less effective than the Wood and Photo versions (ts>3.1; Cohen’s ds>0.195; holm Ps<0.005). The **Question** main effect was also significant (F(3,81)=3.26; P<0.026; η²p=0.108), depicting the fact that the Wonder question produced higher scores than the other three versions (ts>3.01; Cohen’s ds>0.23; holm Ps<0.028).

Examining the 20 Question × Illusion combinations, Wood obtained the highest score in 13 cases (10 following the Wood–Photo–Drawing order). Drawing never ranked highest. Among the remaining seven cases, three involved the Oppel-Kundt illusion and three the Blivet. The distribution of the highest score between the 3 Versions (13-7-0) reached statistical significance (Chi-2 test: Chi²=8.7, p<0.02).

### Ranking analysis

An even clearer pattern emerged from the explicit ranking made by the subjects when shown the three versions of each illusion simultaneously. The Wood version ranked first in all 15 Question × Illusion combinations. The 15 Chi² tests were significant, with values ranging from 15.7 to 68.3 (Ps<0.005). Confirming the previous observation, the 3 Chi² values obtained for the Oppel-Kundt illusion were among the 5 smallest values (15.7 to 25.8). l A logistic regression mixt model indicated that the odds of assigning a higher rank to the Wood illusion were 6.3 times greater (95% HPD CI [4.7, 8.3]) than to the Photo version (see Figure 3., contrasts and structure of the model are detailed in Annex 2).

**Figure 3.**
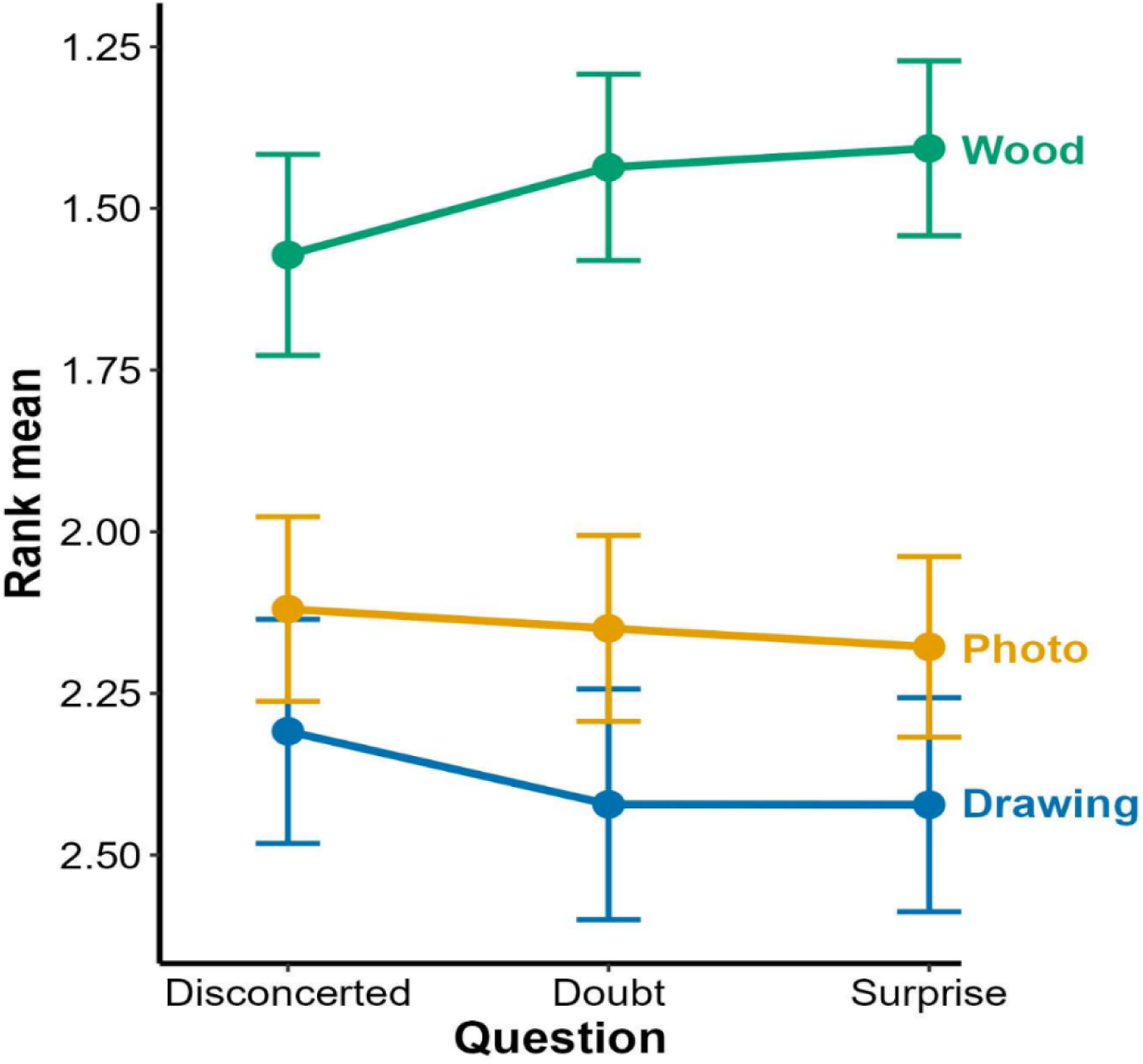
Ranking illusion versions. Ranks ranged from 1 (stronger effects) to 3 (weaker effects). Error bars indicate SEs. Illusion types are color-coded. Ranking of Wood stimuli lied reliably nearest to 1 (i.e. it was most frequently ranked first).

## Discussion

We compared three versions of several illusions to determine whether drawings, photographs, or tangible 3D wooden objects most effectively challenge our perceptual system. Two main conclusions emerged: (1) 3D objects elicited the strongest amazement for impossible objects, and (2) drawings and cognitive illusions had weaker effects, while 2D photographs impact did not exceed 3D objects.

In principle, seeing an impossible object that is simultaneously fully tangible and geometrically impossible should be more challenging to our cognitive system than facing 2D representations because the latter are, or may be, unambiguously artifactual proxies of the real object (Snow and Culham 2021). Our results confirm this prediction. However, slight variability in the 3D viewing angle may have weakened the illusory effects compared with images, which offered a fixed and optimal viewpoint. This is coherent with the observation that the illusion that was most sensitive to the viewing angle, i.e. the Blivet, showed similar results for Wooden and Picture versions (**Annexes** A2). Although this degree of freedom may have reduced the overall impact of 3D illusory effect, our results indicate that the power of the wooden stimuli remained strongest. Although qualitative, the spontaneous reactions of participants are coherent with this quantitative result. In addition, this further extends conclusions drawn by Sugihara and Pinna, 2024) about the power of rectangularity in our 3D construction from 2D information. Not only impossible objects can be constructed, but their paradoxical dimension appears to be most powerful when presented as tangible objects. Provided that our images were embedded in the same 3D environment as the wooden objects, the current study distinguishes between the perception of pictorial objects and tangible objects rather than between 2D and 3D perception.

Our perceptual system can straightforwardly distinguish between possible and impossible objects in 2D (Shacter et al., 1991; Shigemura et al., 2004). According to Ferretti (2024), a key difference between real objects and images lies in access to image boundaries. Thus, interpretation depends not only on intrinsic object geometry but also on contextual cues. Exploration of visual illusions in virtual reality conditions suggests that these cognitive cues may also apply (Škola et al. 2021). The graspable criterion frequently discussed about distinguishing between 2D and 3D stimuli (e.g. Snow and Culham, 2021; Ferretti 2025) suggests that the puzzlement produced by impossible objects may result not only from the violation of geometrical rules but also from their inferred manipulative awkwardness. A physical object, even when only viewed, is graspable and engages sensorimotor brain areas and affords potential actions (grasping, touching, moving around it) in a way that a flat image cannot (Snow and Culham 2021). The impossibility of the object thus creates a deep-seated contradiction with the brain’s action-oriented spatial models. Thus, when violations of geometrical rules and 3D construction converge, they may produce a profound ‘perceptual shock’ that challenges the brain’s core spatial models of reality.

The control, filled-space illusion (Oppel-Kundt; Wakermann, 2017), which relies on cognitive appraisal of distance rather than geometric violation, elicited minimal surprise compared with impossible objects. This suggests that the violation of low-level geometrical rules operates at a deeper encapsulated level than the apprehension of illusions produced by either 2D stimuli or cognitive illusions. Thus, the automatic nature of perceptual 3D construction mechanisms (Sugihara and Pinna 2024) increases the possible dissonance between perceptual processes and geometrical reasoning. By contrast, in the case of tangible impossible objects the resulting conflict is therefore not merely between an image and our knowledge, but between the direct sensory evidence of a "real" object and a lifetime of ingrained knowledge about how the physical world must behave. One may argue that we are processing a tangible perceptual oxymoron rather than a conceptual one.

Further experiments are needed to determine which geometrical features are most likely to generate such perceptual challenge, i.e. difficulties in interpreting and understanding sensory information from the environment. Kokichi Sugihara’s families of 3D illusory objects based on ambiguous cylinders should allow to test specifically for location, orientation, shape, and numerosity effects on our perceptual system (Sugihara 2015a, 2015b, 2018^4^).

Our results may also point to limits for the use of virtual reality and to the need for extensive training in order to develop its realistic power (Harris et al. 2019; Škola et al; 2021, Snow and Culham 2021). Quesque et al. (2023) proposed that visual illusions can be used in order to challenge perceptual evidence (see also Rossetti et al. 2006) and that this may generalize to cognitive levels, which opens new avenues for rehabilitation (Rossetti et al. 2021). We emphasize here that such effects should be optimized by the use of real objects instead of mere images. While images can be engaging, 3D objects confront perception more directly by presenting impossible structures as tangible reality.

Kambara (2017) showed that brief exposure to 2D motion illusions can correct the cognitive bias blind spot, i.e. the tendency to under-evaluate our own susceptibility to cognitive biases. Later, she demonstrated that the effect occurred with paper prints but not with screen-based versions (Kambara, 2021). This attention-grabbing result suggests that although the contrast produced by a computer screen are usually producing more vivid motion illusion, paper versions made it more undisputable to the subjects that motion was illusory. We thus conclude that illusory effects are more powerful when associated with tangible stimuli, i.e. paper vs screen for 2D motion and objects vs images for geometrical illusions. According to our previous proposition to use illusions to develop distance from the evidence of perception and to further develop social cognition, the present results suggest that using tangible stimuli should lead to a stronger impact. We may thus postulate that 3D set-up producing more vivid depth movement illusions should give rise to even stronger cognitive effects.

At a time when most information, real and fake images, are being communicated through 2D interface, it may be reassuring that violation remains most striking when it appears in a natural vs a pictured scene. As we currently encounter a growing body of 2D and ‘3D’ virtual environments and interfaces, and of difficulties disentangling real and fake information and images (e.g. Alexander et al. 2022), and although they might have once appeared trivial, these findings serve as a powerful reminder of the irreplaceable impact of physical experience and offer powerful tools to investigate reality thresholds as well as their pathological alterations (e.g. Keogh 2024, McKormick et al. 2017). They may also provide simple cues and reminders for educational purposes.

### Limitations

Social expectation bias might have affected the explicit ranking of the three types of stimulus presentation. However, the initial scoring phase gave rise to the same pattern of ranking between stimulus versions. Additionally, future studies should investigate the contribution of individual daily screen time to potential alterations of perception.

We tested only four impossible objects and one control illusion. Future studies should examine a broader range of both simple and complex stimuli to assess generalizability. Although we selected rather unfamiliar illusions, one may not exclude that visual illusions being most frequently encountered as images they may in turn appear more surprising when presented as tangible objects. The facts that 1) not only the surprise rating showed a similar pattern, and 2) the cognitive illusion induced the least surprise, indicate that this potential bias should not explain all results.

Looking at objects monocularly through a hole is obviously less ecological than free-viewing conditions (Gibson 1978). Although our image stimuli were embedded in the same 3D environment than the wooden objects, one cannot exclude that motion parallax or accommodation affected perception. The ideal experiment should produce large enough stimuli that can be watched binocularly from far distance, so that they produce two identical retinal images. However, lighting conditions would be more difficult to control under large scale conditions. Therefore, the current study distinguishes between the perception of pictorial objects and tangible objects rather than between 2D and 3D perception.

## Research Transparency Statement

### General Disclosures

#### Artificial intelligence

No artificial intelligence assisted technologies were used in this research or the creation of this article.

#### Ethics

This study was conducted in accordance with the Declaration of Helsinki (World Medical Association, 2013). In compliance with Article R1121-1 of the French Jardé Law, this research did not fall under the category of interventional research involving the human person (RIPH), as it involved healthy volunteers for the purpose of conducting experiments in the field of human and social sciences. This study did not aim to advance biological or medical knowledge.

#### Preregistration no. Materials

All study materials are publicly available in the manuscript, but tangible object cannot.

#### Data

All primary data are publicly available (https://osf.io/p7ywu/files/osfstorage/69655c07c92f7c5b4f5fa346). Analysis scripts: All analysis scripts are publicly available (see annexes).

*Part of this work has been presented at ECVP 2021 : Rossetti, Y; Jeanvoine-Clarac, M; Souilhol, M, Veldeman, M; Volland, F; Haesebaert, F (2021) Learning from illusions : from ambiguity to alterity. Perception 50, 1S, 46-46*.

## Authors contributions

Conceptualization : MS, FV, AK, YR; Methodology: MS, FV, YR; Data Curation : MS, YR; Resources : MS, FV, YR; Software : AF; Formal Analysis: MS, AF, YR; Investigation: MS, YR; Writing – Original Draft Preparation : MS, AF, JE, FQ, YR; Writing – Review & Editing : MS, AF, JE, FQ, FV, AK, YR; Supervision : YR.

## Acknowledgements

The authors wish to thank Sonia Alouche, Laura Lucarelli, and Eric Koun for their dedicated and valuable technical support, Camille Lépingle, Jean-Baptiste Van der Hanst and Frédéric Haesebaert for their comments on an earlier version of this ms. The authors declared that they had no conflicts of interest with respect to their authorship or the publication of this article.

## ANNEXES

**Figure A1:**
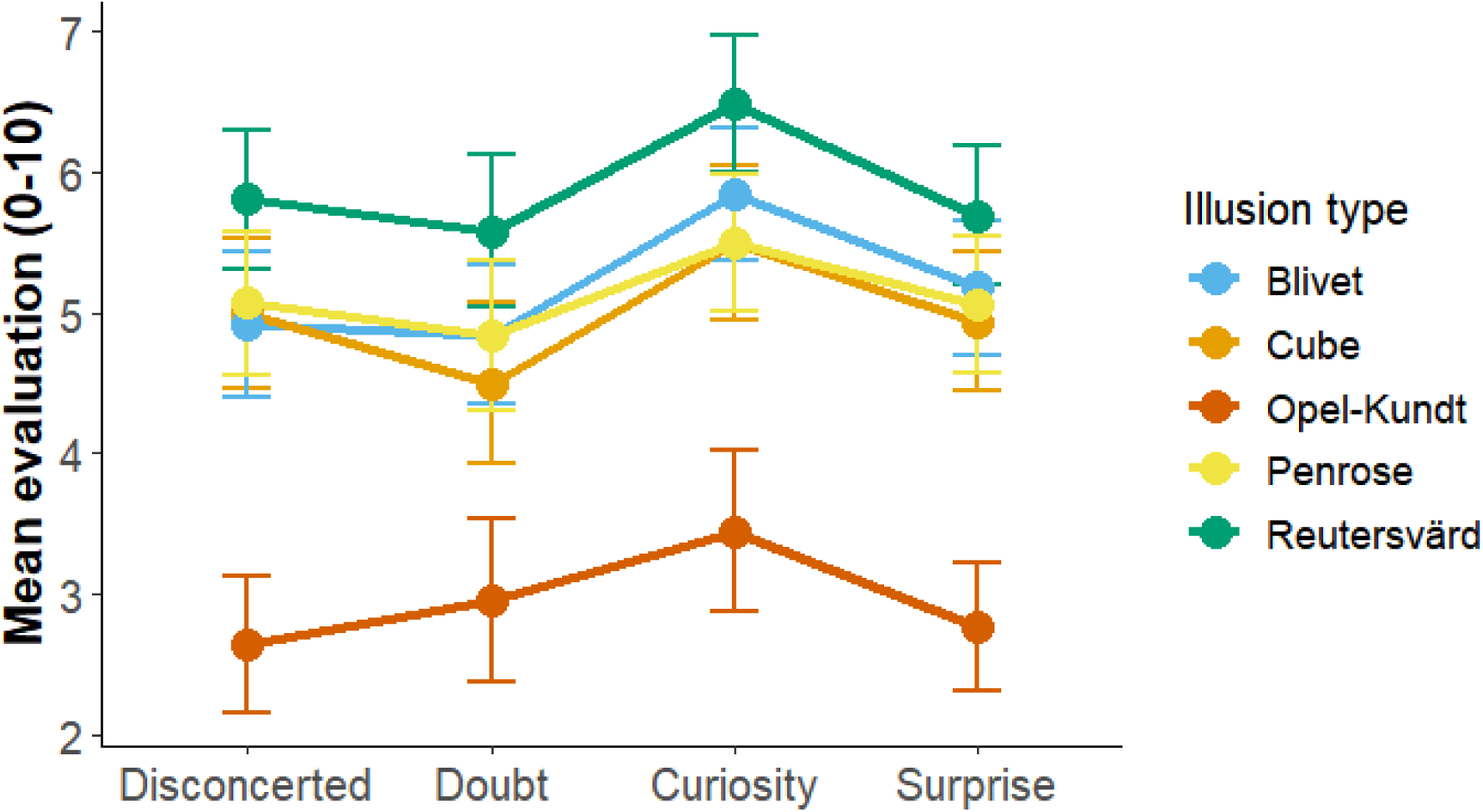
The illusion effect in the rating task. Rating obtained for each illusion, averaged across presentations. The impossible objects showed a homogeneous pattern of results, but the cognitive illusion (Oppel-Kundt) yielded to significantly lower ratings.

**Figure A2:**
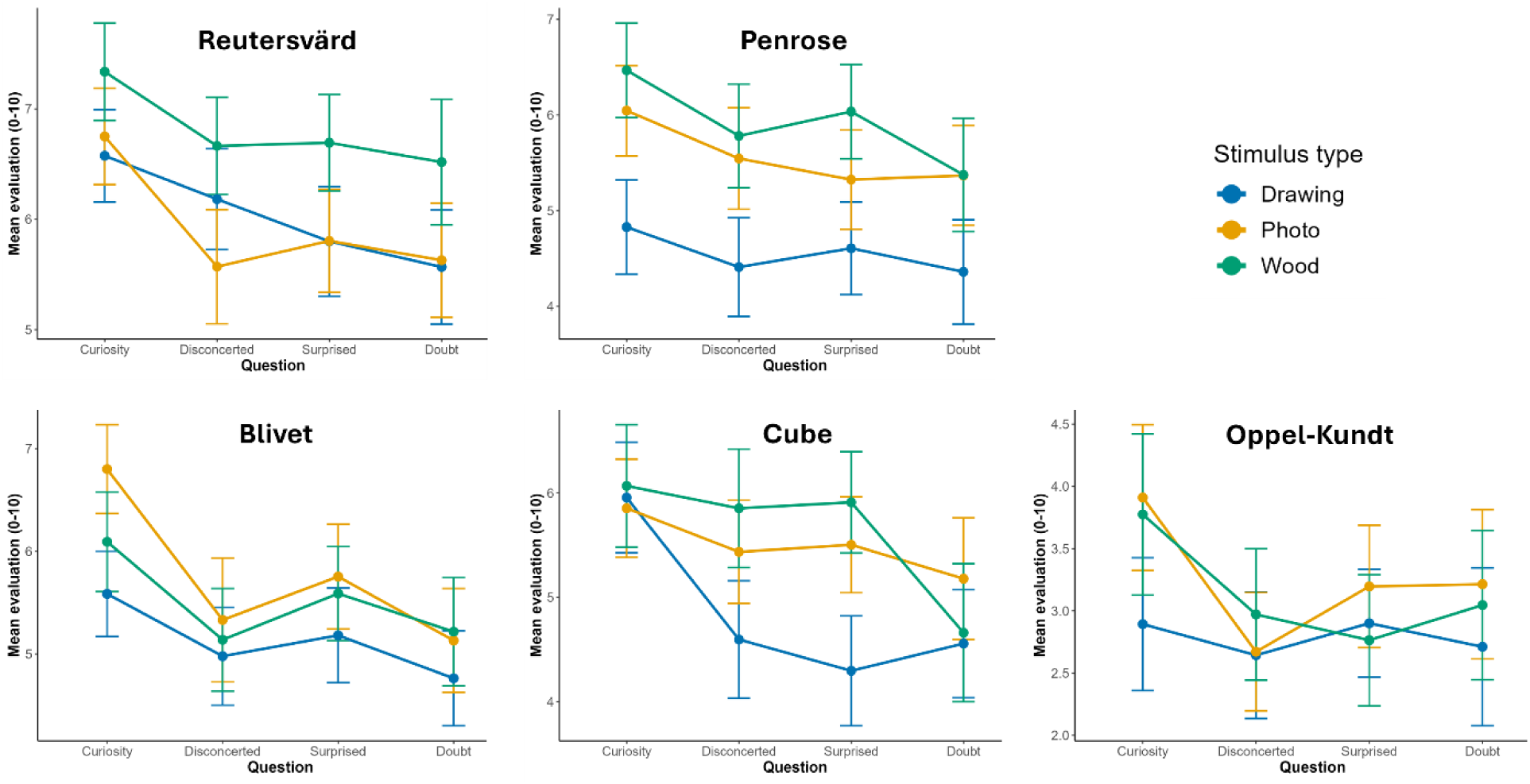
Presentation version effect for each stimulus. Rating obtained for each illusion displayed for each presentation. For the most straightforward impossible objects, i.e. the Reutersvärd frame, the Penrose Triangle and the Cube the Wood presentation raked first for almost all questions. For the cognitive Oppel-Kundt no reliable advantage was found for any version. For the Blivet block, i.e. the impossible object for which the viewing angle was most critical, the Wood version tended to be only higher than the drawing version.

### Annex 1 : Post-Hoc Power Analysis

To justify the sample size, a post-hoc power analysis was conducted for the repeated-measures ANOVA comparing the three stimulus formats (3D wooden object, 2D photograph, and 2D drawing). We used Cohen’s conventions for effect sizes: small (f = 0.10), medium (f = 0.25), and large (f = 0.40):

**Table A1:** Post-hoc power analysis. **Interpretation:** With N = 28 participants, the study has adequate power to detect large effects and reasonable power to detect medium effects, confirming that the sample size is sufficient to identify meaningful differences between the three stimulus formats.

| Effect Size | Power |
| --- | --- |
| Small ($f = 0.10$) | 0.12 |
| Medium ($f = 0.25$) | 0.51 |
| Large ($f = 0.40$) | 0.91 |

The analysis was performed using the pwr package in R (version 4.5.1) with the following script:

~~~
# Install pwr package if necessary
# install.packages("pwr")
library(pwr)
# Repeated-measures ANOVA parameters
k <- 3 # number of conditions
n <- 28 # number of participants
alpha <- 0.05
# Standard Cohen effect sizes
effects <- c(small=0.10, medium=0.25, large=0.40)
# Compute power for each effect size
pwr_results <- sapply(effects, function(f) pwr.anova.test(k=k, n=n, f=f, sig.level=alpha)$power)
~~~

### Annex 2 : Cumulative Logit Mixed-Effects Model Details

The cumulative logit mixed-effects model (CLMM) was fitted to analyze the ordinal ranks assigned to illusions (`value_ordered`, with inverted levels). The model was estimated using Bayesian inference via the `brms` package (Bürkner, 2017), which interfaces with Stan for MCMC sampling.

Below, we provide a detailed description of the model specification, random effects structure, fixed effects, and convergence diagnostics.

#### Model Specification

- **Outcome variable**: value_ordered (ordinal, with inverted levels).
- **Link function**: Cumulative logit (family = cumulative()).
- Formula: `value_ordered ∼ IlluNumb * IlluType + (1 | SUBJECT) + (1 | QuestionClass)`
- Prior distributions:
- Fixed effects: Normal(0, 3)
- Random effects intercepts and standard deviations: Student-t(3, 0, 2.5)

#### MCMC Sampling

- Chains: 2
- Iterations per chain: 5,000 (with 2,500 warmup iterations)
- Thinning: 1
- Total post-warmup draws: 5,000
- Adaptation parameters: adapt_delta = 0.99, max_treedepth = 15
- Seed: 123

#### Random Effects Structure

The model included random intercepts for **subjects (SUBJECT)** and **question classes (QuestionClass)** to account for inter-individual and inter-question variability, respectively. The estimated standard deviations and 95% credible intervals (CrI) for the random effects are summarized in **Table A2**.

#### Fixed Effects and Odds Ratios

The fixed effects estimates (on the log-odds scale) and their corresponding **odds ratios (OR = exp(Estimate))** with 95% highest posterior density intervals (HPD) are presented in **Table A3**. Odds ratios greater than 1 indicate an increased likelihood of assigning a higher rank to the illusion, while odds ratios less than 1 indicate a decreased likelihood.

#### Convergence Diagnostics

Convergence was assessed using **Rhat**, **Bulk_ESS**, and **Tail_ESS** values. All parameters exhibited **Rhat = 1.00**, and effective sample sizes (`Bulk_ESS` and `Tail_ESS`) were greater than 1,000, indicating satisfactory convergence (see Tables B and C).

#### Fixed effects and random effects results in Table B and Table C

**Table A2.**
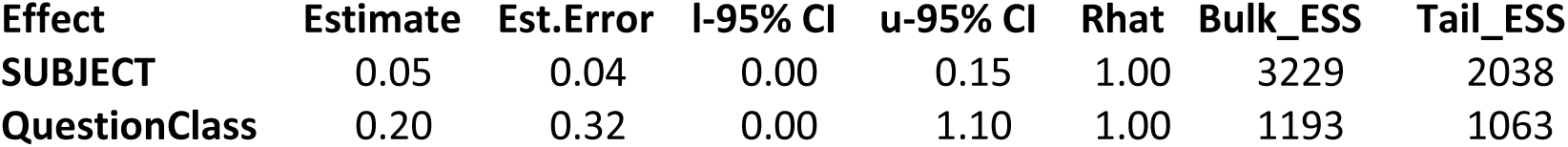
Random Effects.

| Effect | Estimate | Est.Error | I-95% CI | u-95% CI | Rhat | Bulk_ESS | Tail_ESS |
| --- | --- | --- | --- | --- | --- | --- | --- |
| SUBJECT | 0.05 | 0.04 | 0.00 | 0.15 | 1.00 | 3229 | 2038 |
| QuestionClass | 0.20 | 0.32 | 0.00 | 1.10 | 1.00 | 1193 | 1063 |

**Table A3.**
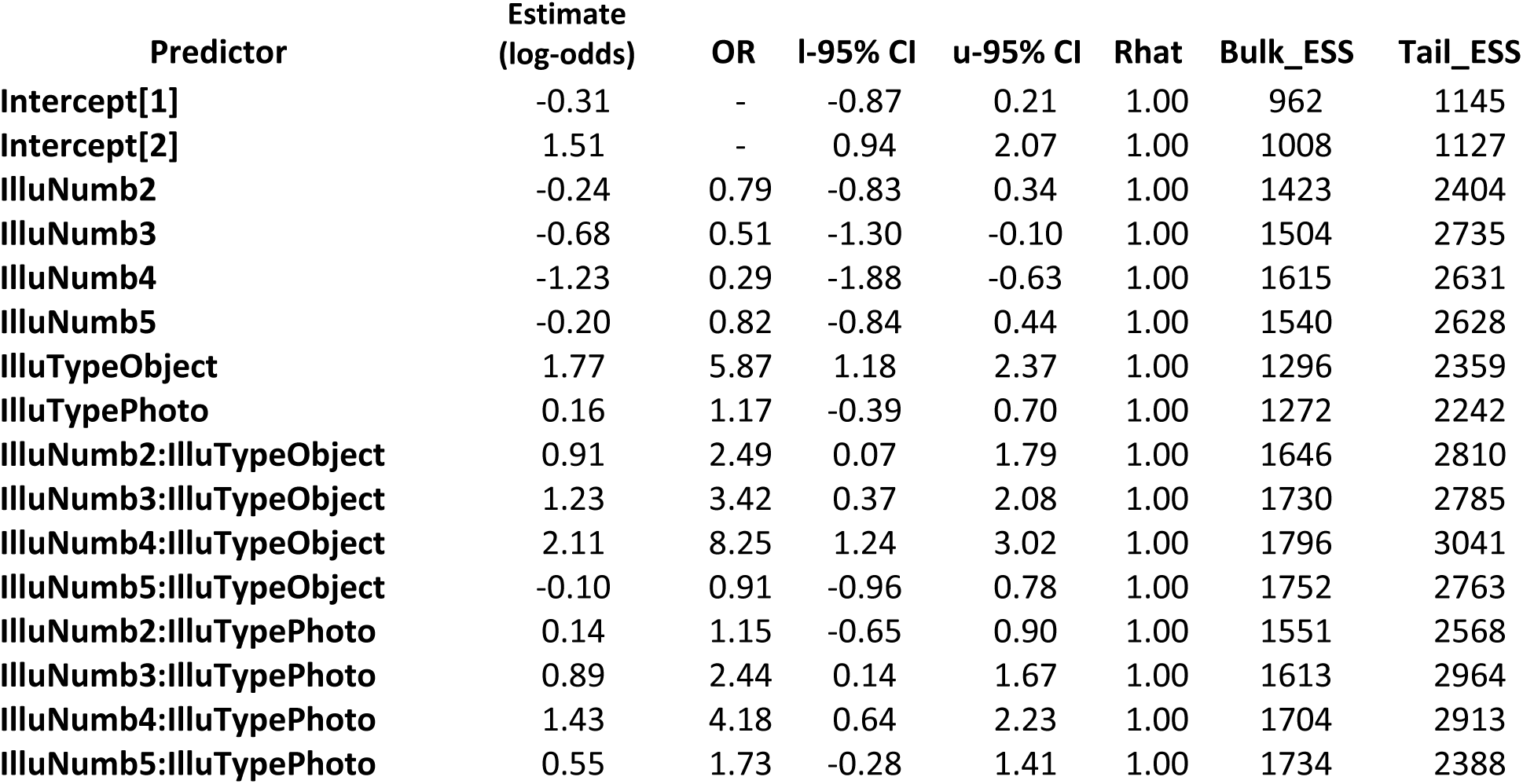
Fixed Effects and Odds Ratios.

| Predictor | Estimate<br>(log-odds) | OR | I-95% CI | u-95% CI | Rhat | Bulk_ESS | Tail_ESS |
| --- | --- | --- | --- | --- | --- | --- | --- |
| **Intercept[1]** | -0.31 | - | -0.87 | 0.21 | 1.00 | 962 | 1145 |
| **Intercept[2]** | 1.51 | - | 0.94 | 2.07 | 1.00 | 1008 | 1127 |
| **IlluNumb2** | -0.24 | 0.79 | -0.83 | 0.34 | 1.00 | 1423 | 2404 |
| **IlluNumb3** | -0.68 | 0.51 | -1.30 | -0.10 | 1.00 | 1504 | 2735 |
| **IlluNumb4** | -1.23 | 0.29 | -1.88 | -0.63 | 1.00 | 1615 | 2631 |
| **IlluNumb5** | -0.20 | 0.82 | -0.84 | 0.44 | 1.00 | 1540 | 2628 |
| **IlluTypeObject** | 1.77 | 5.87 | 1.18 | 2.37 | 1.00 | 1296 | 2359 |
| **IlluTypePhoto** | 0.16 | 1.17 | -0.39 | 0.70 | 1.00 | 1272 | 2242 |
| **IlluNumb2:IlluTypeObject** | 0.91 | 2.49 | 0.07 | 1.79 | 1.00 | 1646 | 2810 |
| **IlluNumb3:IlluTypeObject** | 1.23 | 3.42 | 0.37 | 2.08 | 1.00 | 1730 | 2785 |
| **IlluNumb4:IlluTypeObject** | 2.11 | 8.25 | 1.24 | 3.02 | 1.00 | 1796 | 3041 |
| **IlluNumb5:IlluTypeObject** | -0.10 | 0.91 | -0.96 | 0.78 | 1.00 | 1752 | 2763 |
| **IlluNumb2:IlluTypePhoto** | 0.14 | 1.15 | -0.65 | 0.90 | 1.00 | 1551 | 2568 |
| **IlluNumb3:IlluTypePhoto** | 0.89 | 2.44 | 0.14 | 1.67 | 1.00 | 1613 | 2964 |
| **IlluNumb4:IlluTypePhoto** | 1.43 | 4.18 | 0.64 | 2.23 | 1.00 | 1704 | 2913 |
| **IlluNumb5:IlluTypePhoto** | 0.55 | 1.73 | -0.28 | 1.41 | 1.00 | 1734 | 2388 |

## Footnotes

1 « L’arrivée d’un Train en Gare de La Ciotat » (french)

2 The history of art provides numerous examples of such anonymous and famous concretizations of illusions (e.g. Arcimboldo, “Primavera” (“The spring”), 1573; Utagawa Kuniyoshi, “At first glance he looks fierce but he is a gentle person” or “A face made of people” 1847; Magritte, “La trahison des images” (or “This is not a pipe”, 1929); Dali, “Gala nue regardant la mer qui à 18 mètres apparaît le président Lincoln” 1975; various street-art and anamorphoses,…). In the line of “the father of the impossible figure” Oscar Reutersvärd, probably most artists emphasized puzzling effects of illusory designs and their power to challenge to 3D geometry (e.g. O Reutersvärd, “Perspective Japonaise” series; ME Escher, Relativity 1935; S Fukuda,) over questioning the concept of reality (but see for example Esher’s “Hand with Reflecting Sphere”). However some of this artistic endeavor was specifically intended to cause havoc with perception in order to specifically question the gap between the present reality and its representation (e.g. Magritte’s famous “La trahison des images” 1929 and following versions). The nicknamed “Le saboteur tranquille” (the quiet saboteur) Magritte has been widely acknowledged for his specific ability to implement this philosophical question in the form of a sensory experience (see Snow and Culham 2021).

3 Oscar Reutersvärd’s impossible triangle designs, (e.g. from “Opus 1, n°293”, 1934 to “The Reutersvärd triangle, monolithic version type A” and “type B”, 1949 via “Hommage à Bruno Ernst, perspective japonaise no 262) appeared as early as from 1934 when Penrose & Penrose Triangle was published in1958.

4 https://www.isc.meiji.ac.jp/~kokichis/Welcomee.html

5 Oscar Reutersvärd’s impossible triangle designs, (e.g. from “Opus 1, n°293”, 1934 to “The Reutersvärd triangle, monolithic version type A” and “type B”, 1949 via “Hommage à Bruno Ernst, perspective japonaise no 262) appeared as early as from 1934 when Penrose & Penrose Triangle was published in1958.

